# Assessing the potential ecological functions, ecosystem services and disservices of megaherbivore rewilding in traditional rural wood-pastures

**DOI:** 10.64898/2025.12.11.693666

**Authors:** Marco Davoli, Martina de Benedetto, Flavia Strani, Mario Cipollone, Andrea Lauta, Stefano Focardi

**Affiliations:** Global Mammal Assessment Program, Department of Biology and Biotechnologies, Sapienza University of Rome, 00185 Rome, Italy; Leibniz Institute for Zoo and Wildlife Research, Alfred-Kowalke-Str. 17, 10315, Berlin, Germany; Departamento de Ciencias de La Tierra, Instituto Universitario de Investigación en Ciencias Ambientales deAragón (IUCA), Universidad de Zaragoza, ES-50009 Zaragoza, Spain; Rewilding Apennines, Gioia dei Marsi, Italy; Istituto dei Sistemi Complessi, Sezione di Firenze, Fiorentino, FI, Italy

**Keywords:** Trophic rewilding, megafauna, large mammals, disturbance regimes, semi-feral livestock, herbivory, Italy

## Abstract

Traditional rural wood-pastures are threatened by the loss of wild herbivory and traditional agro-silvo-pastoral practices, promoting woody encroachment, habitat homogenization, and erosion of biodiversity and cultural values. Megaherbivore rewilding has been proposed as a strategy to restore disturbance regimes in these semi-natural landscapes, but its socioecological implications remain uncertain. Using Italy as a representative case study, we reconstructed an early-Holocene, pre-agricultural baseline from fossil records to identify candidate megaherbivores for reintroduction across Alpine, Continental, and Mediterranean biogeographical regions. We compared extant and potential assemblages in functional diversity, vegetation consumption, grazing–browsing balance, and seed-dispersal potential, assessed the sensitivity of conservation-priority habitats to increased herbivory, and reviewed literature through a SWOT analysis of ecosystem services and disservices. The potential assemblage comprised ten species, including extant, range-restricted, new-native, and extinct taxa. Reintroductions would generate uneven functional gains, with limited change in the Alpine region but stronger expansion in the Continental and especially Mediterranean regions, where functional dispersion would increase from 0.46 to 0.64 and vegetation consumption from 4.45 to 10.58 t km^−2^ yr^−1^. Rewilding may shift assemblages towards grazing, potentially helping to limit woody encroachment and reduce fuel loads, but increasing risks for habitats with low-to-moderate herbivory requirements. The SWOT analysis highlighted opportunities for nature-based tourism, rural livelihoods, damage-prevention measures, and integrated wildlife–livestock–human health surveillance, alongside threats related to crop damage, disease transmission, road collisions, management costs, and social conflict. Overall, megaherbivore rewilding should be regarded as a context-dependent restoration strategy requiring spatially explicit planning, density thresholds, and participatory governance.

## Introduction

Wood-pastures, characterized by scattered light-demanding trees within shrubland–grassland mosaics, occur both as naturally maintained open woody ecosystems and as traditional, semi-natural cultural landscapes. In many parts of the world, such as African savannas, wood-pasture-like systems are primarily shaped by natural disturbance regimes, including wild herbivory, fire, and climatic constraints on woody vegetation (Staver *et al*., 2021). By contrast, in landscapes characterized by historical land use such as Europe, wood-pastures depend on the interaction between residual wild herbivory and extensive agro-silvo-pastoral practices, including extensive livestock farming and selective woodland clearing. These landscapes represent key reservoirs of biodiversity and cultural heritage (Moreno et al., 2018; Plieninger *et al*., 2015).

However, in traditional rural wood-pastures both wild and anthropogenic sources of disturbance are being lost. Wild megaherbivores (>45 kg body mass), which play key roles in maintaining vegetation openness and heterogeneity, have undergone long-term declines (Malhi *et al*., 2016). Many species have vanished or been functionally replaced by domestic livestock, while surviving wild populations are increasingly restricted to marginal areas (Davoli *et al*., 2024a; Pacifici *et al*., 2020). For centuries, traditional agro-silvo-pastoral practices partly compensated for this loss by maintaining open mosaic landscapes across much of rural Europe (Giannini and Gabbrielli, 2013; Torralba *et al*., 2018). More recently, rural depopulation and the decline of pastoral economies have reduced livestock activity and vegetation management, promoting woody encroachment, structural simplification, and landscape homogenization. This trend, however, is geographically uneven and is particularly pronounced in Europe and, more broadly, in highly industrialized countries (Kuemmerle *et al*., 2016) (Figure 1).

**Figure 1.**
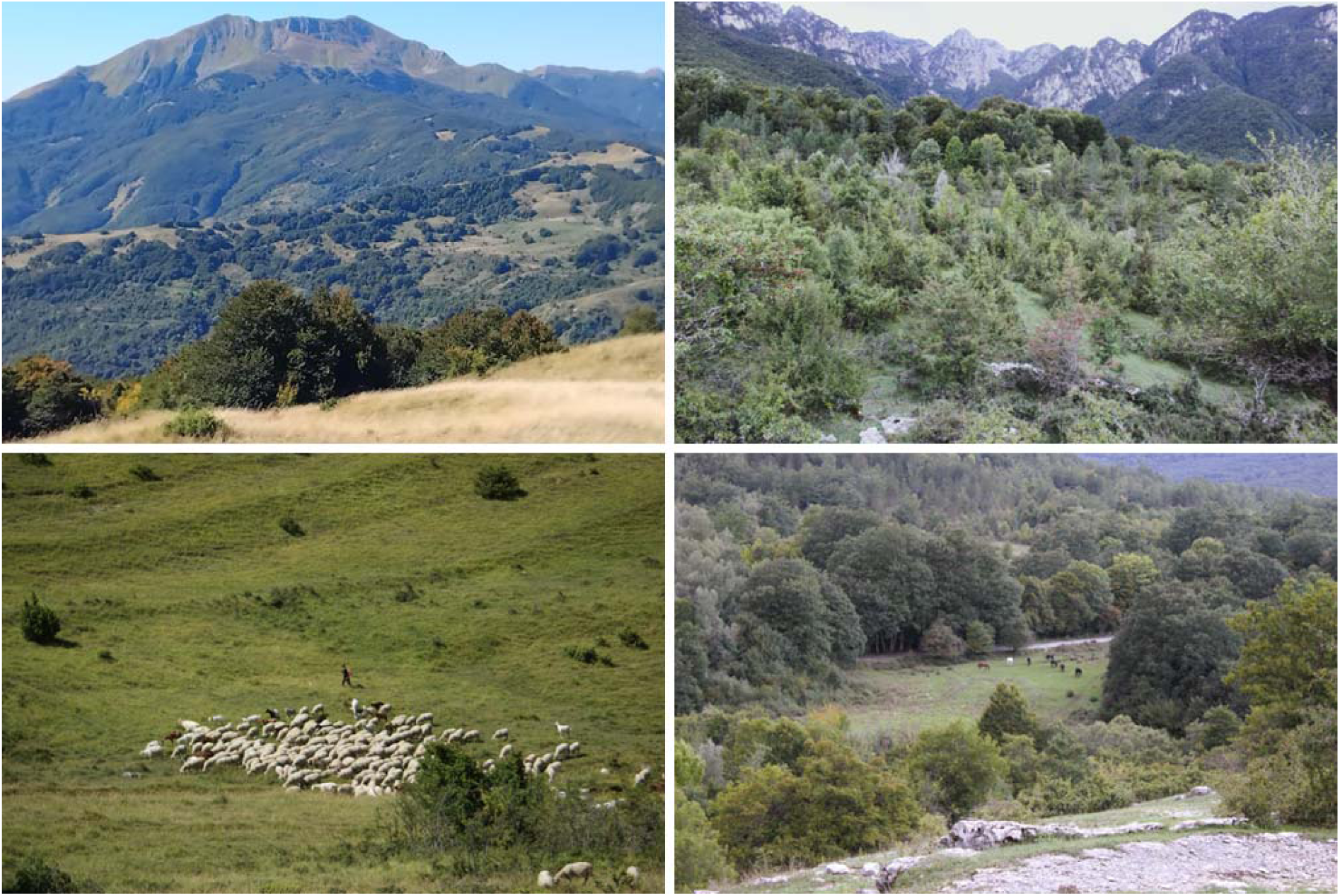
Decline and persistence of semi-wild herbivory in traditional rural wood-pasture systems. Top: landscapes of northern and central Apennines undergoing afforestation following farmland abandonment. Bottom: Free-ranging livestock (sheep, left; horses, right) in central Italy, where traditional agro-silvo*-*pastoral practices still sustain semi-natural mosaics. Photo credits: Marco Davoli, Michela Pacifici.

The restoration of natural dynamics to halt biodiversity decline has emerged as a policy priority withi*n* the European Union (EU), especially following the adoption of the Nature Restoration Law (EU Regulatio*n* 2024/1991) in 2024. Traditional rural wood-pasture systems are recognized for sustaining high habita*t* heterogeneity supporting species-rich communities — for instance, farmland birds (Šálek *et al*., 2025) — while also maintaining the cultural identity of rural people. Importantly, however, the regulation sets onl*y* broad targets and grants considerable flexibility to EU member states regarding implementation. Translatin*g* the aspirations of the Nature Restoration Law into context-specific strategies remains a significant challeng*e* for stakeholders (Hering *et al*., 2023). Notably, current policies still do not explicitly address the historica*l* impoverishment of European megaherbivore communities and the associated decline in key ecologica*l* functions (Garrido *et al*., 2025). Despite this limitation, there is growing recognition among stakeholders tha*t* contemporary landscapes lack herbivory disturbance historically pivotal to the evolution of biodiversity. I*n* response, process-based approaches such as rewilding have gained momentum, supported by growin*g* public interest in restoring nature (Bergin *et al*., 2024; Jones and Comfort, 2020). However, assessing th*e* relevance of megaherbivore rewilding for wood-pasture systems restoration first requires identifying whic*h* species could plausibly constitute a functionally meaningful reference assemblage for reintroductions.

Unlike conventional restoration, rewilding emphasizes the reestablishment of trophic complexity by facilitating the return of functionally important species (Mutillod *et al*., 2024; du Toit and Pettorelli, 2019). Within this framework, megaherbivore restoration seeks to enhance top-down regulation on vegetation structure and resource dynamics (Wang *et al*., 2023). This is particularly relevant for Europe, where the marked decline in megafaunal diversity since the Late Pleistocene has led to ecological dynamics that diverge from evolutionary conditions (Davoli *et al*., 2024b). Furthermore, the loss of grazing and browsing disturbance favours the expansion of shade-tolerant plant communities less characteristic of wood-pasture systems, while also increasing fuel loads (Czyżewski and Svenning, 2025; Corlett, 2025; Kirkland *et al*., 2024). However, the extent to which reintroduced megaherbivores would complement or merely replicate the ecological functions currently provided by extant assemblages remains unclear. Likewise, identifying which habitats are most likely to benefit from restored herbivory — and which may instead be vulnerable to potential negative impacts — is essential for assessing the conservation implications of megaherbivore rewilding.

Rewilding offers opportunities to revitalize depopulated rural areas by promoting multifunctional landscapes that generate socioeconomic benefits (Garcia-Ruiz *et al*., 2020; Perino *et al*., 2019). These may include enhanced ecotourism, strengthened local identity, greater community engagement in natural resource management, and notably, hunting opportunities (Hall, 2019; Massenberg *et al*., 2023). However, the adoption of rewilding strategies remains constrained by persistent uncertainties, especially regarding their alignment with stakeholder interests and the capacity to yield socially acceptable outcomes (Cary *et al*., 2025). Additionally, reintroducing megaherbivores into human-dominated landscapes poses major practical challenges and requires careful, context-specific evaluation. Key concerns include potential ecosystem disservices, such as livestock depredation by recovering large carnivores supported by renewed prey bases (Recio *et al*., 2020), crop damage (Hearn *et al*., 2014), unintended effects on disease dynamics (Rafael et al., 2026), and other real or perceived risks. Moreover, because contemporary landscapes are shaped by altered species assemblages and modified disturbance regimes, rewilding may produce nonlinear ecological outcomes rather than simply restoring functions consistent with evolutionary baselines (Hoeks *et al*., 2023). Further uncertainty concerns species selection and ecological equivalence, as illustrated by debates over whether extinct taxa should be functionally replaced by extant relatives, such as the wisent as a proxy for the steppe bison formerly present in the Iberian Peninsula (Nores *et al*., 2024; but see Bartolomé *et al*. 2026 for an alternative interpretation). Thus, evaluating megaherbivore rewilding as a landscape-management strategy requires assessing not only its potential to restore ecological functions, but also the implications for rural communities, including potential ecosystem services and disservices.

Building on these considerations, this paper critically evaluates the socioecological implications of megaherbivore rewilding, considering the impact of potential reintroductions anchored to a pre-agricultural, Holocene baseline (approximately the last 12,000 years). Italy provides a compelling context because it still harbours a relatively diverse set of wild megaherbivores and long-standing cultural landscapes shaped by millennia of interactions between natural dynamics and traditional land use, such as transhumant pastoralism (Troiano *et al*., 2021), while also having experienced severe megaherbivore range contractions or extirpations over recent centuries (Palombo *et al*., 2024). Specifically, this study addresses the following research questions: (1) What would constitute the potential megaherbivore assemblage using a pre-agriculture, Holocene reference? (2) Would reintroduced species be functionally redundant with extant assemblages? (3) Which habitats stand to benefit, or potentially be threatened, by megaherbivore rewilding? (4) What are anticipated implications, including both ecosystem services and disservices, for rural communities?

## Material and methods

### Overview

Using fossil records, we reconstructed the Italian megaherbivore assemblage that occurred after the last glacial period and before major land-use changes associated with the advent of agriculture and farming. This provided a Holocene, pre-agricultural ecological baseline for identifying species with potential reintroduction relevance in Italy based on nativeness, thereby addressing Research Question 1. Species were classified across Italy’s three biogeographical regions, which are distinct environmental and management units: Alpine, Continental, and Mediterranean. This subnational framework avoids extrapolating reintroduction potential to the entire peninsula for species ecologically restricted to specific areas.

For each biogeographical region, we independently compared the extant assemblage, comprising species currently present, with a potential assemblage that additionally included regionally extirpated species. We then quantified changes in ecological functions to address Research Question 2. To address Research Question 3, we assessed which habitat types listed in Annex I of the Habitats Directive occur in Italy and how they are distributed across the three biogeographical regions. Based on the expected increase in vegetation consumption under the potential assemblage, we qualitatively identified habitats that could either benefit from or be negatively affected by megaherbivore rewilding. Finally, to address Research Question 4, we conducted a ‘Strengths, Weaknesses, Opportunities, Threats’ (SWOT) analysis to evaluate the broader ecosystem services and disservices potentially associated with megaherbivore rewilding in Italian rural areas.

### Megaherbivore assemblages and functional traits

We reviewed literature to gather evidences of megaherbivore occurrences in Italy between 12,000 and 6,000 years ago, corresponding to the early-middle Holocene interglacial phases and predating the Neolithic (i.e., agricultural revolution) (Biagi and Spataro, 2002; Skeates, 2003). Our fossil records dataset was primarily based on Mondanaro *et al*. (2025) and supplemented by additional bibliographic sources, retaining only temporally calibrated records supported by standard dating methods (Table S1). For the extant assemblage, we compiled an inventory based on Loy *et al*. (2019), potentially including new native species (sensu Lemoine and Svenning, 2022): natural migrants (colonised within the last 500 years without human assistance) and historic introductions (human-assisted arrivals between 10,000 and 500 years ago). To reach the potential assemblage, we added to the extant assemblage all regionally extirpated species with at least one unequivocal Holocene fossil record.

For each considered species, we then collected functional traits including body mass (log_10_-scaled), digestive system type, limb morphology, and the relative dietary importance of graminoids (grazing) and browse/fruits (browsing), as reported in Lundgren *et al*. (2021) (Table S2). We also extracted species density under natural conditions (individuals km^-2^) and estimated potential vegetation consumption as dry matter intake (t km^−2^ yr^−1^) from Pedersen *et al*. (2023), based on phylogenetically adjusted allometric models of population density in core areas of extant populations and metabolic demand. Further methodological details and assessments of estimate reliability are provided therein.

### Ecological functions of herbivore assemblages

We quantified and compared the functional diversity (FD) of the herbivore guild in the extant and potential assemblages. We expected FD to increase only where reintroduced species in a biogeographical region may potentially contribute novel functional traits not represented within the extant assemblage. FD enables comparisons of community-level functional differences by characterizing species across multiple ecologically relevant traits, which are transformed into orthogonal axes (via principal component analysis - PCoA) to account for trait covariance (Laliberte and Legendre, 2010; Villéger *et al*., 2008). FD is determined by the geometry of multidimensional trait space, particularly the convex hull defined by species’ trait combinations. Functionally distinct species that extend this convex hull increase FD, reflecting greater functional complementarity (Mouillot *et al*., 2013). Notably, we quantified FD using Functional Dispersion (FDis), scaled between 0 and 1, which measures the abundance-weighted mean distance of species to the assemblage centroid in functional trait space. For trait space construction we used all the functional traits collected. Moreover, we used potential densities under natural conditions to generate abundance weights for functional traits (at a spatial resolution of 1 km², corresponding to the scale of the potential density data). Functional indices were computed using the *mFD package* in R *v3.4.1* (Magneville *et al*., 2022). For simplicity, and given the macroecological comparative scope of the study, we retained only the first two PCoA axes, which captured most of the functional trait variation. Notably, the *mFD package* allows functional traits to be weighted by species’ relative densities, ensuring that more abundant species exert a proportionally greater influence on FD. This abundance-weighted approach provides a closer approximation of ecological reality by reflecting the unequal contribution of species to community-level functional processes.

Furthermore, we quantified key ecological effects of the extant and potential assemblages, including total potential vegetation consumption, the relative contribution of grazing versus browsing, and mean daily travel distance (as a proxy for seed-dispersal potential) as follows: 1) potential vegetation consumption per km² was obtained by summing consumption values for all species in each assemblage, following the approach of Davoli *et al*. (2024b). 2) The grazing – browsing balance of each assemblage was quantified using a community-level diet proportion index (DP), weighted by species’ body mass. For each species, we combined body mass with proportions of graminoid and browse/fruit consumption to compute a diet score ranging from 0 (entirely graminoid-based) to 1 (entirely browse/fruit-based), which was then aggregated across the assemblage. The elaborated formula is:

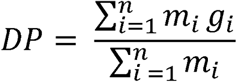

where *m* represents body mass, *g* denotes the importance of graminoids as a proportion of the diet, and *n* represents the total number of species. Notably, graminoid importance data in Lundgren *et al*. (2021) are reported as an ordinal variable with the categories 0 (0–9% of the diet), 1 (10-24%), 2 (25-49%) and 3 (50–100%). To enable quantitative trait analyses, we converted these ordinal classes into continuous fractional values. Specifically, for each ordinal category we assigned three alternative values corresponding to the minimum, mean, and maximum percentage of the diet represented by the category’s defined range (e.g., for 50–100%: min 50, mean 75, max 100). This approach allowed us to account for uncertainty in dietary composition while preserving the original categorical structure of the data. 3) Seed dispersal potential was estimated as the modelled mean daily Euclidean travel distance (D, km day⁻¹) using the energetic/search framework of Carbone *et al*. (2005), in which D scales with body mass (M, kg) to the ¼ power:

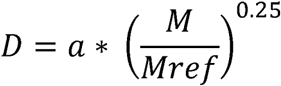

The calibration constant a was derived from field data on red deer (*Cervus elaphus*), selected for its intermediate body size within our potential assemblage and its ability to persist in human-modified landscapes while remaining relatively elusive. It was set to 2.88 km day⁻¹ based on telemetry data from a semi-natural landscape in the Massif Central, France, broadly comparable to Italian rural conditions (Pépin *et al*., 2009).

### Light and herbivory requirements of priority habitats

We compiled a comprehensive list of conservation-priority habitats listed in Annex I of the Habitats Directive that occur in Italy, using the ‘Italian Interpretation Manual of the Habitats Directive’ (Biondi *et al*., 2010). We restricted the analysis to habitats classified as natural grassland formations, semi-natural dry grasslands and shrub-covered facies, temperate European forests, Mediterranean deciduous forests, temperate montane coniferous forests, and Mediterranean montane coniferous forests (Table S3), assuming that abiotic dynamics predominantly governed disturbance regimes in all other habitats, such as coastal systems. We retained only conservation-priority habitats marked with an asterisk in the Directive as requiring priority action at the EU level (EUR-Lex 92/43/EEC).

For each selected habitat, we identified the diagnostic plant taxa representative of the Italian context, as outlined in Biondi *et al*. (2010). For each taxon, we assigned quantitative indices that describe the realized niche optima for light and herbivory requirements. Specifically, we used ecological indicator values (light requirement) and disturbance indicator values (herbivory disturbance requirement) compiled by Dengler *et al*. (2023) and Midolo *et al*. (2023) for native European plants. These indices, calculated as weighted averages of environmental conditions across species’ observed occurrences, approximate plant taxa’s optimal realized niches. As relative measures, they do not quantify the absolute herbivory pressure required by each habitat, but indicate which habitats are more likely to benefit from, or be negatively affected by, increased herbivory.

We categorized the indicator values into discrete classes of light and herbivory requirement. Light requirement values ranged continuously from 0 to 10 in the database, whereas herbivory requirement ranged from 0 to 1. We assigned categories relative to the empirical distribution of all values in the database, i.e., across all European plant species considered. We defined four classes: *low* (values below the first quartile), *mid-low* (between the first quartile and the median), *mid-high* (between the median and the third quartile), and *high* (values above the third quartile). Lastly, using the most recently available national habitat mapping for Italy produced by the Italian Institute for Environmental Protection and Research (ISPRA Report 2013-2018), we qualitatively assessed the spatial distribution and potential vulnerability of these habitats across the three biogeographical regions. Each habitat was classified as absent, rare, or widespread, and as likely to benefit from, be threatened by, or respond context-dependently to increased herbivory.

### SWOT analysis of ecosystem services and disservices

We conducted a systematic literature review to support a SWOT analysis (Ghazinoory *et al*., 2011) aimed at evaluating potential services and disservices of megaherbivore rewilding in rural Italy, similarly to Treves *et al*. (2020). Search for the literature review was performed in Web of Science (last accessed on 11 June 2026), limited to open-access, peer-reviewed articles in English published between 2021 and 2026. We restricted the review to this period as a trade-off between obtaining sufficiently comprehensive evidence base and maintaining the feasibility of the analysis, given that each selected study required examination and extraction of relevant information. A search string was developed using the TS field to automatically screen titles, abstracts, and keywords. The query combined terms referring broadly to medium- and large-sized mammals potentially targeted by population restoration interventions, as well as species commonly regarded as problematic or damaging in conservation and management contexts. For instance, we considered also large carnivores because their responses to restored wild prey bases may indirectly generate ecosystem services and disservices. The search string further incorporated terms linked to restoration ecology and rewilding, while explicitly excluding aquatic systems and saprophytic organisms, and applied geographic filters to Italy and its administrative regions: “TS = (mammal* OR megafauna* OR megaherbivore* OR megacarnivore* OR herbivore* OR carnivore*) AND TS = (rewild* OR “restor*” OR introduc* OR reintroduc* OR rebound* OR return OR feral OR free-ranging) NOT TS = (fish* OR marine OR sea OR saproph*) AND TS=(Italy OR Italian OR Apennines OR “Italian Alp*” OR Abruzzo OR Basilicata OR Calabria OR Campania OR Emilia-Romagna OR Friuli-Venezia Giulia OR Lazio OR Liguria OR Lombardia OR Lombardy OR Marche OR Molise OR Piemonte OR Piedmont OR Puglia OR Apulia OR Sardegna OR Sardinia OR Sicilia OR Sicily OR Toscana OR Tuscany OR Trentino-Alto Adige OR South Tyrol OR Alto Adige OR Trentino OR Umbria OR Valle d’Aosta OR Aosta Valley OR Veneto)”.

All retrieved articles were screened at the title and abstract level by two independent reviewers (with cross-checking and a required double-positive decision) to exclude irrelevant studies. The final set included only publications that explicitly addressed the expected direct implications of megaherbivore rewilding for people in contemporary Italian rural landscapes. From each eligible study, summary statements relevant to the SWOT framework were extracted and assigned to one of four predefined dimensions: (i) Strengths — intrinsic positive outcomes of rewilding for ecosystems and society (e.g., habitat restoration, enhanced ecosystem services). (ii) Weaknesses — internal limitations or risks (e.g., management complexity, unintended ecological impacts). (iii) Opportunities — external drivers facilitating rewilding (e.g., land abandonment, growing environmental awareness, policies, eco-tourism). (iv) Threats — external pressures or adverse consequences (e.g., human–wildlife conflict, zoonotic risks, political instrumentalization). Individual statements were then grouped by thematic similarity into overarching macro-statements, which were used to generate the summary SWOT matrix.

## Results

### Potential megaherbivore assemblages and reintroduction candidates

We compiled 91 georeferenced and chronologically constrained records from 18 fossil sites dated to 11.4–6 ka across the Italian Peninsula, including four multilayered sites (Figure 2; Table S1). Of the six extant megaherbivores, ibex (*Capra ibex*) and brown bear (*Ursus arctos*) have restricted contemporary ranges relative to their historical distributions and were therefore treated as regional reintroduction candidates where currently absent but historically present. Two species, fallow deer (*Dama dama*) and mouflon (*Ovis aries orientalis*), are new natives sensu Lemoine and Svenning (2022). Four additional taxa from the fossil baseline are regionally or globally extinct in the wild: aurochs (*Bos primigenius*), European wild ass (*Equus hydruntinus*), moose (*Alces alces*), and wild horse (*Equus ferus*). Together, these taxa define a potential assemblage of ten species (Figure 2). Under fossil-inferred biogeographical constraints, the potential assemblage includes brown bear, fallow deer, mouflon, ibex, red deer, wild boar (*Sus scrofa*), and the candidate reintroductions aurochs and moose in the Alpine region; fallow deer, mouflon, red deer, wild boar, and the candidate reintroductions aurochs, brown bear and ibex in the Continental region; fallow deer, mouflon, red deer, wild boar, and the candidate reintroductions aurochs, European wild ass, ibex, and wild horse in the Mediterranean region.

**Figure 2.**
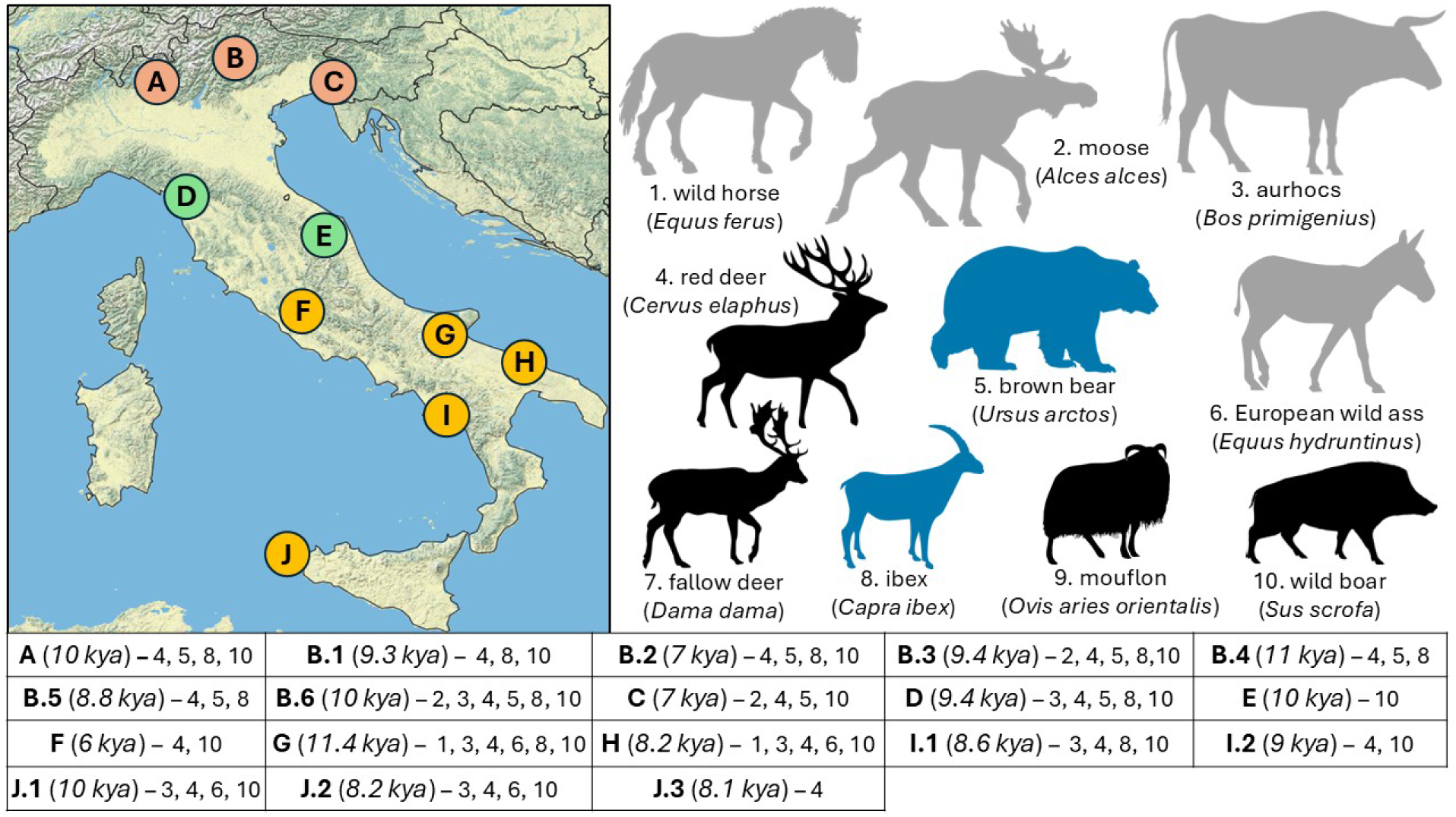
Fossil sites and potential megaherbivore assemblage. Top left: fossil sites from Mondanaro *et al*. (2025), grouped by biogeographical region: brown = Alpine, green = Continental, orange = Mediterranean. Top right: potential megaherbivore assemblage in Italy, ordered by increasing body mass from bottom left to top right. Black indicates species in the extant assemblage; blue indicates extant species with markedly reduced ranges, which were therefore considered candidates for reintroduction in some regions; grey indicates species extinct in the wild (grey). Bottom: fossil sites are labelled with alphanumeric codes corresponding to Table S1, with approximate ages in kya and identified species, following the numbering in the assemblage panel. Map lines delineate study areas and do not necessarily depict accepted national boundaries.

### Ecological effects of extant vs potential megafauna assemblages

Across the three biogeographical regions, PC1 was primarily associated with variation in browsing diet, whereas PC2 mainly reflected differences in digestive fermentation type (both p < 0.05). The largest increase in functional dispersion between the extant and potential assemblages would occur in the Mediterranean (FDis: 0.46 extant vs 0.64 potential; Figure 3-1), whereas the change would be more limited in the Alpine (FDis: 0.50 vs 0.56). The potential reintroduction of brown bear in the Continental and wild horse in the Mediterranean would substantially expand the functional trait hypervolume of these regional assemblages (Figure 3-2).

**Figure 3.**
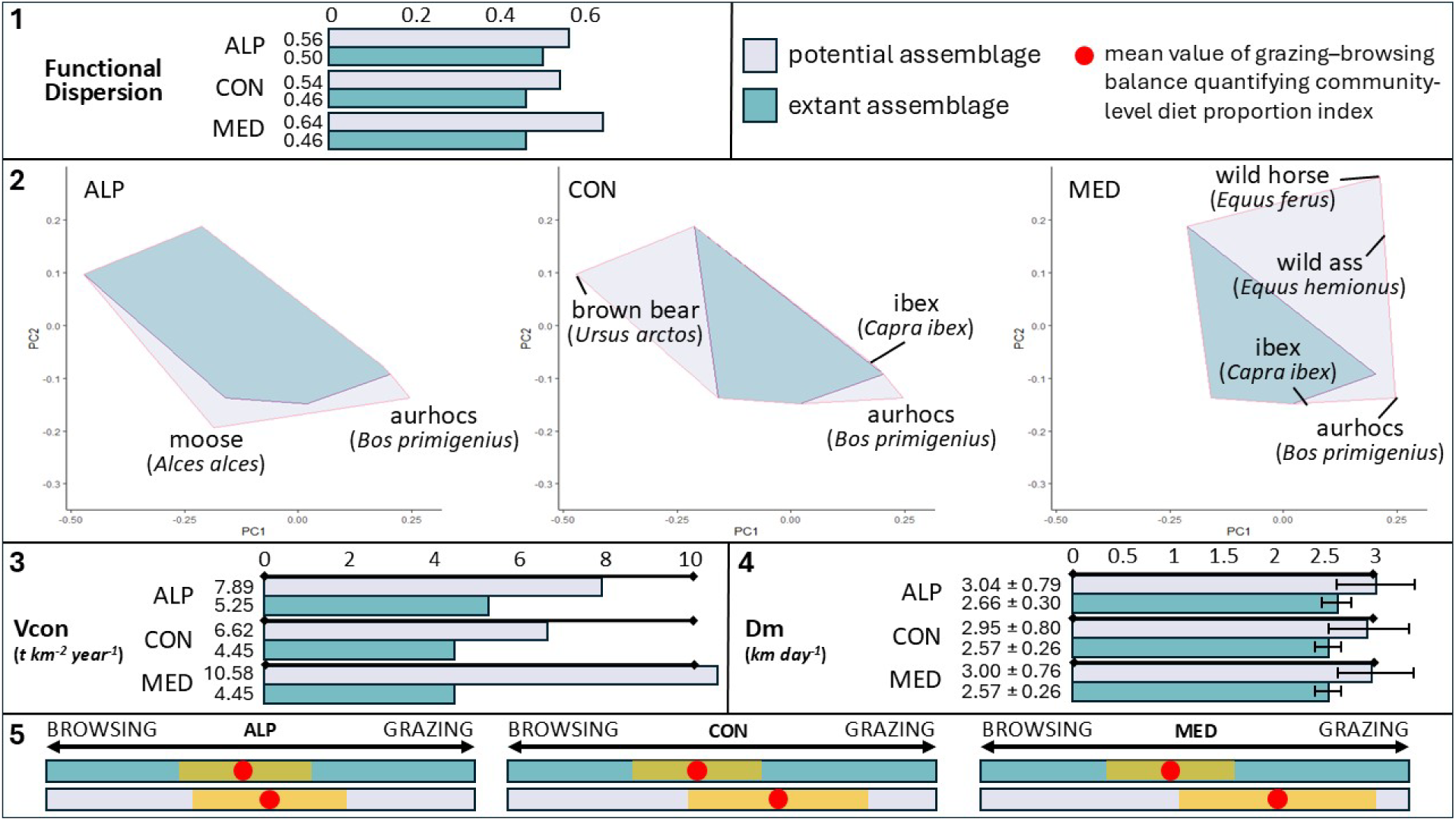
Comparison of potential ecological functions between the extant and potential megaherbivore assemblages in the three biogeographical regions. (1) Comparison of functional dispersion; (2) functional diversity hypervolumes (PC1 and PC2); (3) Comparison of estimated vegetation consumption (Vcon, t km⁻² yr⁻¹); (4) seed-dispersal potential approximated by the Euclidean distance of daily movements (Dm, km); (5) Comparison of potential browsing vs grazing herbivory prevalence.

Similarly, the largest potential increase in vegetation consumption would be in the Mediterranean biogeographical region, rising from 4.45 t km⁻² yr⁻¹ in the extant assemblage to 10.58 t km⁻² yr⁻¹ in the potential assemblage (Figure 3-3). Mean daily movement distance would be broadly similar between extant and potential assemblages across the three biogeographical regions (Figure 3-4).

**Figure 4.**
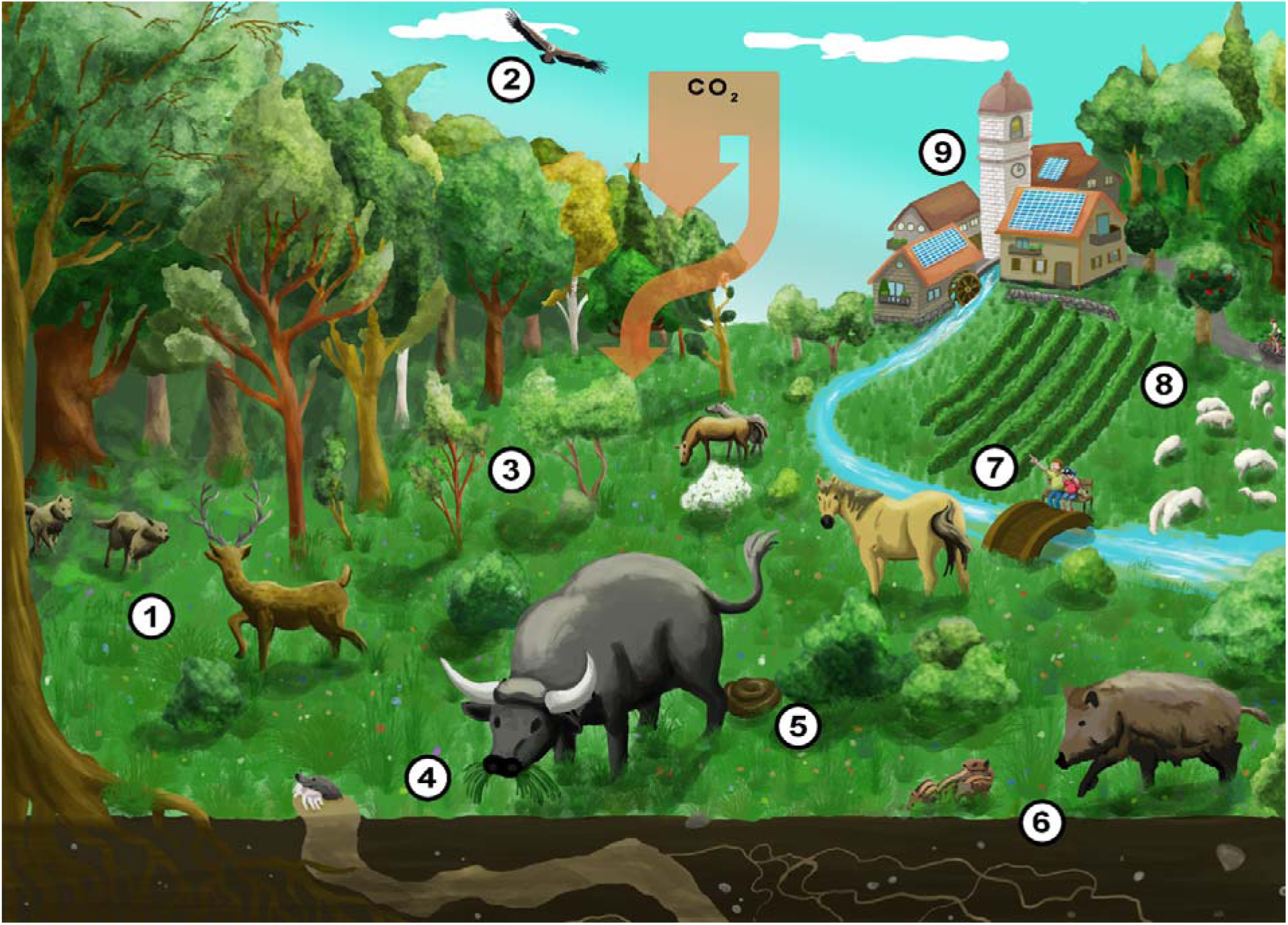
Potential ecosystem services and disservices following megaherbivore rewilding in traditional rural wood-pasture systems. Numbers indicate: 1) positive and negative effects of large carnivore recover*y* under enhanced prey availability; 2) cascading effects across multiple trophic levels; 3) altered vegetation*-*use patterns; 4) increased overall vegetation consumption; 5) increased dung deposition and nutrien*t* redistribution; 6) increased soil disturbance; 7) enhanced interactions between people and nature; 8) renewed integration of extensive livestock systems; and 9) closer integration of rural communities withi*n* more nature-rich landscapes.

The proposed reintroductions would shift the diet of megaherbivore communities from predominantly browsing- to predominantly grazing-oriented assemblages. This shift would be particularly marked in the Mediterranean biogeographical region, where the mean ratio would increase from 0.44 ± 0.14 in the extant assemblage to 0.69 ± 0.22 in the potential assemblage, on a scale ranging from 0, indicating a fully browsing assemblage, to 1, indicating a fully grazing assemblage (Figure 3-5).

### Potential impact on conservation-priority habitats

Conservation-priority habitats were classified for light requirement into four categories: low (< 6.90), mid-low (6.90–7.69), mid-high (7.70–8.39), and high (≥ 8.40). Similarly, habitats were classified for herbivory requirement as low (< 0.194), mid-low (0.194–0.242), mid-high (0.243–0.308), and high (≥ 0.309). Accordingly, one open habitat (‘62.60’) showed high requirement for both light and herbivory disturbance. Three habitats (two open and one forestal: ‘61.10’, ‘62.40’, ‘95.60’) showed mid-high requirements, while two open habitats (‘62.10’, ‘62.30’) showed mid-low light requirement but mid-high herbivory disturbance requirement. All other habitats were associated to low or mid-low requirements. Of the 17 habitats considered, megaherbivore rewilding would be expected to have a positive impact on four, with two additional habitats potentially to benefit from increased grazing but not requiring high light levels. For all remaining habitats, expected impact of megaherbivore rewilding could be negative or mildly negative (Table 1).

**Table 1.** Expected impact of megaherbivore rewilding on conservation-priority habitats. For each priority EUNIS-coded habitat occurring in Italy, we assessed its distribution across biogeographical regions and estimated mean light and herbivory-disturbance requirements from diagnostic plant taxa. Requirements are expressed on relative scales of 0–10 and 0–1, respectively. Expected effects were qualitatively classified assuming increased herbivory in landscapes where each habitat occurs.

| Habitat type | Light requirement | Grazing pressure | Distribution and qualitative assessment of expected impact |
| --- | --- | --- | --- |
| 61.10 Rupicolous calcareous or basophilic grasslands of the *Alysso-Sedion albi* | mid-high<br>(8.29 ± 0.45) | high<br>(.33 ± .02) | Widespread in Continental and Mediterranean regions. <u>Expected positive impact.</u> |
| 62.10 Semi-natural dry grasslands and scrubland facies on calcareous substrates | mid-low<br>(7.56 ± 0.53) | mid-high<br>(.28 ± .03) | Widespread everywhere. <u>Expected positive impact with enhanced grazing.</u> |
| 62.20 Pseudo-steppe with grasses and annuals of the *Thero-Brachypodietea* | high<br>(8.49 ± 0.73) | high<br>(.36 ± .05) | Widespread in Continental and Mediterranean regions. <u>Expected positive impact.</u> |
| 62.30 Species-rich *Nardus* grasslands, on siliceous substrates in mountain areas | mid-low<br>(7.28 ± 0.61) | mid-high<br>(.27 ± .03) | Widespread in Continental and Mediterranean regions. <u>Expected positive impact with enhanced grazing.</u> |
| 62.40 Sub-pannonic steppic grassland | mid-high<br>(8.09 ± 0.63) | high<br>(.33 ± .03) | Rare and only in Alpine region. <u>Expected positive impact.</u> |
| 91.80 Tilio-Acerion forests of slopes, screes and ravines | low<br>(4.14 ± 1.05) | low<br>(.19 ± .01) | Widespread in Alpine region but rare elsewhere. <u>Expected negative impact.</u> |
| 91.AA Eastern white oak woods | low<br>(5.94 ± 0.97) | mid-low<br>(.23 ± .02) | Widespread in Continental and Mediterranean regions. <u>Expected negative impact.</u> |
| 91.D0 Bog woodland | low<br>(6.48 ± 1.41) | low<br>(.19 ± .04) | Rare and only in Alpine region. <u>Expected negative impact.</u> |
| 91.E0 Alluvial forests with *Alnus glutinosa* and *Fraxinus excelsior* | low<br>(5.75 ± 0.69) | low<br>(.18 ± .02) | Widespread in Alpine and Continental regions. <u>Expected negative impact.</u> |
| 91.H0 Pannonian woods with *Quercus pubescens* | low<br>(6.44 ± 0.84) | mid-low<br>(.23 ± .02) | Widespread in Alpine region and rare in Continental region. <u>Expected negative impact.</u> |
| 92.10 Apeninne beech forests with *Taxus* and *Ilex* | low<br>(3.55 ± 1.08) | low<br>(.19 ± .01) | Widespread in Mediterranean region and rare in Continental region. <u>Expected negative impact.</u> |
| 92.20 Apennine beech forests with *Abies alba* and beech forests with *Abies nebrodensis* | low<br>(3.92 ± 1.55) | mid-low<br>(.20 ± .00) | Rare and only in Continental and Mediterranean regions. <u>Expected negative impact.</u> |
| 94.30 Subalpine and montane *Pinus uncinata* forests | mid-low<br>(7.16 ± 0.61) | mid-low<br>(.24 ± .02) | Rare and only in Alpine and Continental regions. <u>Expected mild negative impact.</u> |
| 95.10 Southern Apennine *Abies alba* forests | low<br>(3.17 ± 1.25) | low<br>(.19 ± .00) | Rare and only in Mediterranean region. <u>Expected negative impact.</u> |
| 95.30 (Sub-) Mediterranean pine forests with endemic black pines | mid-low<br>(7.21 ± 1.10) | mid-low<br>(.24 ± .06) | Rare and only in Alpine and Mediterranean regions. <u>Expected mild negative impact.</u> |
| 95.60 Endemic forests with *Juniperus* spp. | mid-high<br>(8.20 ± 0.60) | high<br>(.31 ± .04) | Rare and only in Alpine and Mediterranean regions. <u>Expected positive impact.</u> |
| 95.80 Mediterranean *Taxus baccata* woods | low<br>(4.69 ± 1.63) | mid-low<br>(.21 ± .02) | Rare and only in Mediterranean region. <u>Expected negative impact.</u> |

### Ecosystem services and disservices of megaherbivore rewilding

Of the 101 papers retrieved from the Web of Science database, 34 (33.7%) were retained after screening. The retained papers were dominated by studies on large carnivores, particularly wolves (including *C. l. italicus*), followed by work on wild boar, red fox (*Vulpes vulpes*), brown bear, livestock systems, beavers (*Castor fiber*), and non-native or invasive mammals relevant to human–wildlife interactions (Table S4). Based on the included papers, we derived 29 summary macro-statements describing potential ecosystem services, disservices, management constraints, and governance opportunities associated with megafauna rewilding in Italy (Table 2). Overall, the synthesis was dominated by disease-related and conflict-related issues. The most frequently addressed opportunities concerned integrated wildlife–livestock–human health surveillance (41.2% of retained papers) and damage-prevention or compensation measures (23.5%), while the most common threats were pathogen and parasite circulation (41.2%), carnivore depredation on livestock (23.5%), and social opposition or polarization around wildlife recovery (17.6%). By contrast, direct evidence for positive ecosystem services to people was comparatively limited, with individual strength macro-statements each supported by no more than 5.9% of retained papers.

**Table 2.**
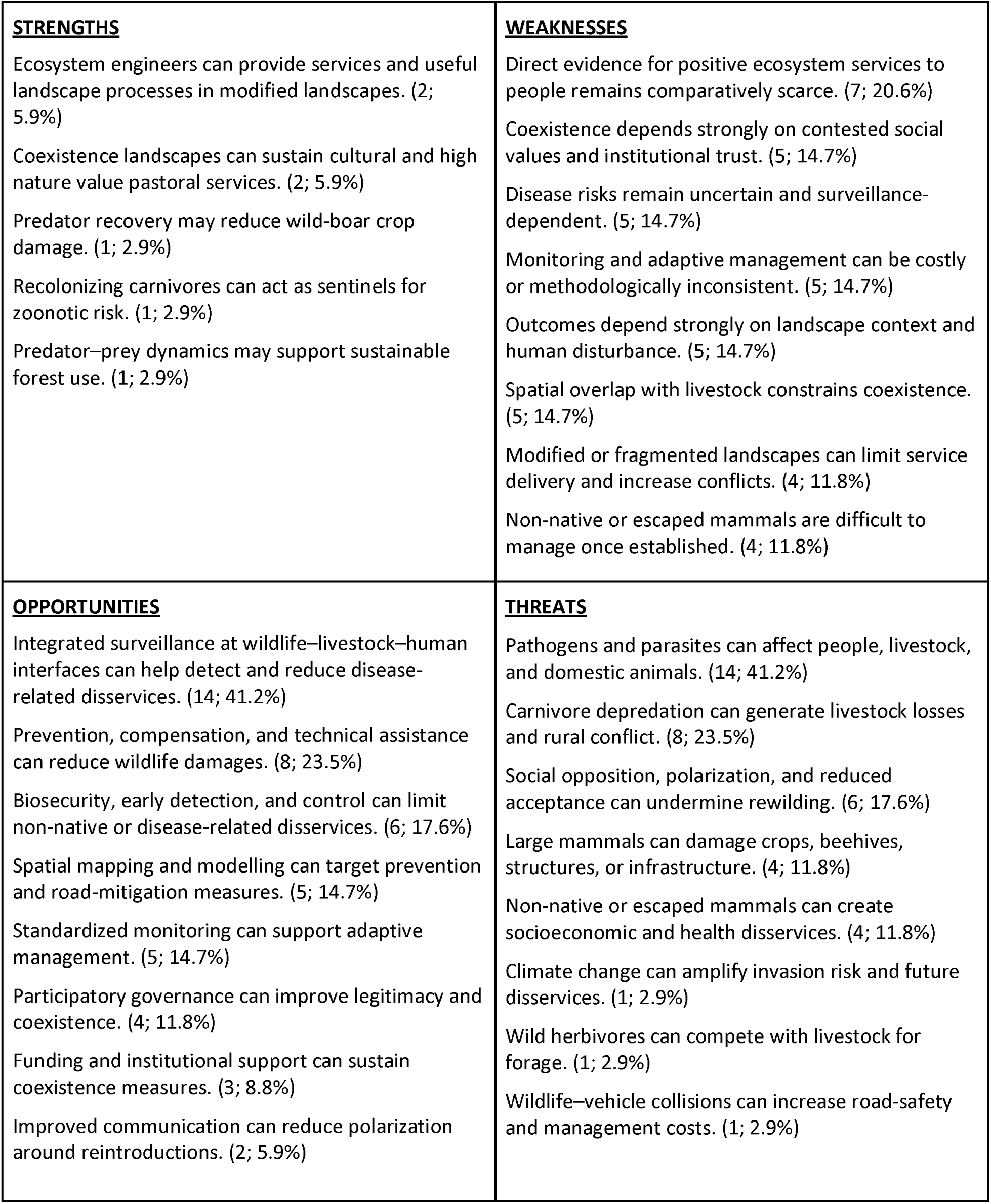
Megaherbivore rewilding and implications for people. SWOT macro-statements based on the screened papers listed in Table S4. Values in parentheses indicate the number of articles and the percentage of retained papers addressing each topic.

|  |  |
| --- | --- |
| **<u>STRENGTHS</u>**<br>Ecosystem engineers can provide services and useful landscape processes in modified landscapes. (2; 5.9%)<br>Coexistence landscapes can sustain cultural and high nature value pastoral services. (2; 5.9%)<br>Predator recovery may reduce wild-boar crop damage. (1; 2.9%)<br>Recolonizing carnivores can act as sentinels for zoonotic risk. (1; 2.9%)<br>Predator–prey dynamics may support sustainable forest use. (1; 2.9%) | **<u>WEAKNESSES</u>**<br>Direct evidence for positive ecosystem services to people remains comparatively scarce. (7; 20.6%)<br>Coexistence depends strongly on contested social values and institutional trust. (5; 14.7%)<br>Disease risks remain uncertain and surveillance-dependent. (5; 14.7%)<br>Monitoring and adaptive management can be costly or methodologically inconsistent. (5; 14.7%)<br>Outcomes depend strongly on landscape context and human disturbance. (5; 14.7%)<br>Spatial overlap with livestock constrains coexistence. (5; 14.7%)<br>Modified or fragmented landscapes can limit service delivery and increase conflicts. (4; 11.8%)<br>Non-native or escaped mammals are difficult to manage once established. (4; 11.8%) |
| **<u>OPPORTUNITIES</u>**<br>Integrated surveillance at wildlife–livestock–human interfaces can help detect and reduce disease-related disservices. (14; 41.2%)<br>Prevention, compensation, and technical assistance can reduce wildlife damages. (8; 23.5%)<br>Biosecurity, early detection, and control can limit non-native or disease-related disservices. (6; 17.6%)<br>Spatial mapping and modelling can target prevention and road-mitigation measures. (5; 14.7%)<br>Standardized monitoring can support adaptive management. (5; 14.7%)<br>Participatory governance can improve legitimacy and coexistence. (4; 11.8%)<br>Funding and institutional support can sustain coexistence measures. (3; 8.8%)<br>Improved communication can reduce polarization around reintroductions. (2; 5.9%) | **<u>THREATS</u>**<br>Pathogens and parasites can affect people, livestock, and domestic animals. (14; 41.2%)<br>Carnivore depredation can generate livestock losses and rural conflict. (8; 23.5%)<br>Social opposition, polarization, and reduced acceptance can undermine rewilding. (6; 17.6%)<br>Large mammals can damage crops, beehives, structures, or infrastructure. (4; 11.8%)<br>Non-native or escaped mammals can create socioeconomic and health disservices. (4; 11.8%)<br>Climate change can amplify invasion risk and future disservices. (1; 2.9%)<br>Wild herbivores can compete with livestock for forage. (1; 2.9%)<br>Wildlife–vehicle collisions can increase road-safety and management costs. (1; 2.9%) |

## Discussion

We provide an assessment of rewilding potential by evaluating how herbivore reintroductions in Italy, grounded in a concept of nativeness informed by a deep-time ecological baseline, could alter ecological functions and generate ecosystem services and disservices. We define the ecological baseline as the early Holocene, after the last glaciation but before large-scale anthropogenic transformations of natural landscape patterns (Kaplan *et al*., 2009). This period reflects faunal assemblages closely aligned with contemporary climatic conditions – although already being substantially impoverished relative to the Late Pleistocene (Sandom *et al*., 2014).

Megaherbivore reintroductions, as a form of trophic rewilding sensu Svenning *et al*. (2016), represent an innovative yet still controversial approach. Restoring diversified herbivory regimes may be particularly relevant for traditional rural wood-pasture systems, whose semi-open habitat mosaics support biodiversity and form part of the cultural heritage. These mosaics are increasingly homogenized by woody encroachment, which can also occur through passive rewilding, with negative consequences for biodiversity (Garrido *et al*., 2021; Bugalho *et al*., 2026). However, megaherbivore rewilding should not be assumed to benefit biodiversity in every ecological context. First, reintroductions should prioritize native species or, where these are extinct, functionally analogous taxa, as species introduced into ecosystems where they did not evolve may generate greater ecological costs than benefits (Bescond-Michel *et al*., 2025). Accordingly, the selection of species for reintroduction should account for their native occurrence within the relevant biogeographical region. Furthermore, rebounding wildlife may generate ecosystem disservices to people, including crop and forestry damage, disease transmission, competition with livestock or new-native herbivores, road collisions, management costs, and social conflicts (Carpio *et al*., 2021; König *et al*., 2020). The outcomes of megaherbivore rewilding are therefore likely to depend on population density, regional context and management regime, with effects that are positive at low or moderate densities potentially becoming negative under excessive population densities (Pringle *et al*., 2023; Trepel *et al*., 2024).

In addressing the potential megaherbivore assemblage for Italy, fossil evidence reveals that four wild megaherbivores — aurochs, European wild ass, moose, and wild horse — were present in Italy during the early-to-mid Holocene. Currently, these species are extinct in the wild or exist only as domestic descendants. Moose vanished from the Alpine region due to habitat fragmentation driven by climatic warming and increased humidity (Marcott *et al*., 2013), with human hunting further accelerating their decline (Schmölcke and Zachos, 2005). Therefore, moose should be considered for reintroduction exclusively within the Alpine biogeographical area. The aurochs, ancestor of domestic cattle and the largest-bodied Holocene megaherbivore recorded in Italy, was widespread across the peninsula, suggesting it could potentially be reintroduced throughout the country. In contrast, the wild horse and European wild ass are known only from sites in the Mediterranean biogeographical region, although both species were widely distributed across Eurasia during the Late Pleistocene, likely due to their dietary flexibility (Boulbes and Van Asperen, 2019; Strani and DeMiguel, 2023). Competition with emerging domestic livestock may have contributed to the extinction of both wild horse and European wild ass, paralleling current interactions observed between Przewalski’s horses and free-ranging domestic ungulates (Sietses *et al*., 2009).

Regarding extant taxa, fallow deer was widespread across Europe, including Italy, during warm Late Pleistocene phases but disappeared by the Last Glacial Maximum (MIS 2), persisting only in the Balkans and Anatolia into the Early Holocene. Zooarchaeological and genetic evidence indicates that humans later translocated fallow deer from the Balkans to western Mediterranean, starting in the Bronze age and particularly during the Roman period, including to Italy and Sardinia (Baker *et al*., 2024a, 2024b; Focardi *et al*., 2025). Similarly, the mouflon was reintroduced to Italy only in the 19^th^ century (Apollonio *et al*., 2010), despite evidence from Eemian-age fossils (cc. 125,000 years ago) indicate that this species was present in Italy in the Late Pleistocene (Davoli *et al*., 2024b). Two megaherbivores persist today but with ranges smaller than their historical distributions: the brown bear and the Alpine ibex. Brown bears suffered severe declines between the 19^th^ and 20^th^ centuries due to persecution and habitat loss, leaving only a remnant alpine population and the critically reduced Apennine subspecies (*U. a. marsicanus*) (Ciucci *et al*., 2017). The brown bear could be considered for reintroduction in Italy’s Continental biogeographical region, where substantial suitable habitat likely remains available, particularly in areas such as the northern Apennines. The Alpine ibex experienced an earlier bottleneck, surviving only in the Gran Paradiso National Park in the early 19^th^ century; although now recovering across the Alps, all modern populations descend from this enclave and retain low genetic diversity (Bozzuto *et al*., 2019). Fossil evidence indicates that this species also occurred in the Continental and Mediterranean biogeographical regions, in addition to the Alpine region.

The results of our analysis of functional redundancy and novelty following reintroductions are not unexpected, as species additions can increase the probability that the functional space occupied by an ecological community will expand. However, such expansion does not arise from species richness per se, but from the incorporation of distinct functional attributes (Dalerum *et al*., 2012), which is precisely the rationale underpinning the reintroduction framework proposed here. This rationale produced contrasting outcomes across the three biogeographical regions considered, suggesting that the ecological justification for rewilding may vary substantially across Italy. In the Alpine region, the proposed reintroductions would generate only limited increases in FD, indicating that the balance between ecological benefits and the considerable practical and social challenges associated with reintroductions may be less favourable. These challenges include not only conflicts between animals and people, but also conflicts among stakeholder groups over conservation priorities and management responsibilities (Beckmann and Soorae, 2022; Deli *et al*., 2023). By contrast, stronger functional gains would be expected in the Continental region through the addition of the large-bodied mixed-feeder brown bear, and especially in the Mediterranean region, where the reintroduction or bovids and equids would substantially increase FD.

The prominent role of bovids and equids is particularly significant, as the domestic descendant of the taxa are typical to extensive livestock systems in traditional rural wood-pastures, thereby partially maintaining herbivory patterns akin to those once produced by their wild ancestors. This functional continuity underscores the potential value of ‘rewilding-lite’ strategies, whereby semi-domestic herbivores are managed in semi-wild, low-intervention settings to restore herbivory, trampling, dung deposition, and other disturbance regimes (Gordon *et al*., 2021; van Dooren *et al.,* 2025). However, policy frameworks remain ambiguous regarding the recognition of semi-wild livestock as trophic replacements for extinct megaherbivores, while how behavioural and morphological differences between wild species and their domestic descendants shape ecological outcomes remains poorly understood (Pérez-Barberìa *et al*., 2023; Mutillod *et al*., 2025). The use of functional analogues well suited to novel ecosystems (Svenning *et al*., 2024), and potentially more acceptable to local communities, may further reinforce this approach. For example, the reintroduction of semi-wild water buffalo (*Bubalus bubalis*) in wetlands, as already implemented in the Danube Delta (Rewilding Europe), may also support culturally relevant ecosystem services, given the importance of buffalo milk and derived products in Mediterranean food traditions.

A particularly relevant finding is that, in the Mediterranean region, the proposed reintroductions would more than double vegetation consumption. This estimate, however, is necessarily sensitive to actual species abundances and their relative proportions, which in our case were assumed from expected natural-density conditions as estimated by Pedersen *et al*. (2023). Through woody encroachment, fuel loads increase drastically and, as a natural consequence, so does wildfire risk (Kirkland *et al*., 2024) — a growing concern for EU policies (Hermoso *et al*., 2022). Critically, this dynamic creates a reinforcing feedback loop at the global scale: higher fuel loads intensify wildfires, wildfires contribute substantially to greenhouse gas emissions, and the resulting climatic warming further heightens wildfire probability (Bowman *et al*., 2009). Moreover, although wildfires can function as abiotic disturbances that maintain biodiversity (Pausas et al., 2025; Viljur *et al*., 2022), unregulated fire frequency and severity may degrade soil properties and, together with other extreme disturbance events, substantially reduce nutrient availability (Morán-Ordóñez *et al*., 2020; Pausas *et al*., 2008).

Restoring wild herbivory could therefore be pivotal in limiting the accumulation of fire-prone woody biomass in traditional wood-pastures, especially because increased grazing has been associated with reductions in wildfire frequency, intensity, flame height, and combustion temperature (Holdo *et al*., 2007; Johnson *et al.,* 2018). In this sense, megaherbivore rewilding could contribute to broader EU restoration objectives aimed at enhancing ecosystem resilience, mitigating climate-related risks, and restoring natural processes in degraded landscapes (Garrido *et al*., 2025). Beyond fire-risk mitigation, enhanced herbivory may also influence carbon cycling by transferring above-ground biomass to soils through dung deposition, potentially supporting mineral-associated organic matter formation and long-term soil carbon sequestration (Kristensen *et al*., 2021; orange arrow in Figure 4). Such below-ground carbon storage can further improve soil structure, water retention, nutrient availability, and long-term soil fertility (Lal *et al*., 2015).

By contrast, our results suggest only limited changes in potential seed dispersal relative to the extant assemblage. Megaherbivores are effective agents of endozoochorous dispersal and reduced megaherbivore assemblages are associated with diminished landscape connectivity and regeneration potential (Fricke *et al*., 2025), as well as a lower capacity for plant communities to track environmental changes (Fricke *et al*., 2022). The limited differences between extant and potential assemblages likely reflect the already impoverished nature of the candidate fauna relative to Plio-Pleistocene assemblages, when European plant communities interacted with a broader range of large-bodied dispersers, including proboscideans associated with plants such as horse chestnut (*Aesculus hippocastanum*) (Tighe, 2026).

Across all three biogeographical regions, the proposed reintroductions would produce a marked shift towards grazing. Although increased grazing does not imply reduced browsing pressure in our analysis, indirect effects mediated by interspecific competition and altered space use cannot be excluded. Such shifts in herbivory regimes can substantially affect vegetation dynamics, particularly in forest understories (Bernes *et al*., 2018), and may therefore have contrasting implications for conservation-priority habitats. In light-demanding habitats such as 61.10*, 62.20*, and 62.40*, increased grazer dominance could reduce woody-plant control, increase shading, and shift communities away from optimal niche conditions, especially in the Mediterranean region where 61.10* and 62.20* are widespread. Conversely, 62.10* grasslands, which are widespread in Italy and linked to traditional mountain pastoral systems, may benefit from increased wild herbivory or the restoration of extensive semi-wild livestock husbandry (Olmeda *et al*., 2019; Filibeck *et al*., 2019). However, because 11 of the 17 conservation-priority habitats assessed showed medium-to-low herbivory requirements, increased herbivore pressure could also generate negative niche shifts, highlighting the need to consider the potential risk of exposing these habitats to excessive herbivory disturbance.

In terms of ecosystem services and disservices, our SWOT analysis suggests that megaherbivor*e* rewilding may generate potential benefits for people, but also management constraints and social or health*-*related risks that cannot be assumed to be straightforwardly resolved (Figure 4). By increasing wild pre*y* availability (Figure 4-1), megaherbivore rewilding may support large-carnivore recovery and potentiall*y* reduce carnivore reliance on domestic animals, thereby lowering livestock depredation risk in some context*s* (Ripple *et al*., 2014; Alquinta *et al*., 2026). However, carnivore recovery may also increase perceived risk, ideological opposition, and conflict with pastoral communities, especially where livestock losses are frequen*t* or prevention measures are insufficient (von Hohenberg and Hager, 2022). Cascading effects across multipl*e* trophic levels (Figure 4-2) may contribute to regulating herbivore impacts, supporting landscap*e* heterogeneity, and indirectly benefiting pollination or other nature-based services, yet these outcomes ar*e* highly context-dependent and may also shift herbivore pressure toward crops, forestry resources, o*r* sensitive habitats (Garrido *et al*., 2022; Davoli *et al*., 2022; Bernes *et al*., 2018). Similarly, altered vegetation*-*use patterns (Figure 4-3) and increased overall vegetation consumption (Figure 4-4) may help limit woody encroachment, maintain open habitats of cultural and conservation value, and reduce fuel loads, but only within appropriate density thresholds; excessive browsing or grazing could instead damage priority vegetation, compete with livestock forage, or reduce local tolerance toward rewilded populations (Côté *et al*., 2004; Garrido *et al*., 2021). Increased dung deposition and nutrient redistribution (Figure 4-5) may enhance nutrient cycling, secondary seed dispersal, and resources for dung-associated biodiversity, while concentrated deposition may create sanitary concerns, nutrient hotspots, and higher parasite-transmission risk at wildlife-livestock-human interfaces (Kristensen *et al*., 2021; Veldhuis *et al*., 2018). Moderate soil disturbance (Figure 4-6) may generate recruitment microsites and maintain fine-scale habitat heterogeneity, whereas intense trampling can compact soils, reduce infiltration, increase runoff and erosion, and facilitate ruderal or alien plant establishment, with potential consequences for pasture quality and land management (Eldridge and Soliveres, 2023; Centeri, 2022; Santoianni *et al*., 2024).

More frequent interactions between people and nature (Figure 4-7) may foster environmental education, public engagement, and nature-based tourism, but may also increase human-wildlife conflict through disease concerns and wildlife-vehicle collisions, thereby undermining social acceptance (König *et al*., 2020; Balčiauskas *et al*., 2025). Renewed integration of extensive livestock systems (Figure 4-8) could align rewilding with traditional agro-silvo-pastoral practices, support rural livelihoods, and maintain grazing-dependent cultural landscapes, but may also intensify wildlife-livestock interfaces involving forage competition, pathogen circulation, and the need for sustained compensation, prevention, and surveillance measures (Barroso and Gortázar, 2024). Finally, closer integration of rural communities within more nature-rich landscapes (Figure 4-9) can support place-based restoration, shared stewardship, and long-term coexistence, provided that governance is participatory and local benefits are visible. Conversely, resistance may increase where costs are unevenly distributed, decision-making is perceived as externally imposed, or communities experience reduced autonomy over land-use practices (Faure *et al*., 2024).

Importantly, several limitations of this study warrant consideration. First, functional analyses assumed a uniform distribution of megaherbivores, a necessary simplification that may obscure spatial heterogeneity in functional effects. However, this assumption was applied consistently to both assemblages, allowing a macroscale comparison focused on relative rather than absolute differences in functional outcomes. Second, estimates of ecological functions were derived from published trait values without explicitly incorporating uncertainty, which may affect precision but does not bias the comparative assessment between assemblages. In addition, the SWOT analysis relied exclusively on academic literature. While this approach enabled a synthesis of dominant research perspectives, it does not capture the views of local stakeholders who directly experience the implications of ongoing wildlife rebounding (Deinet *et al*., 2013). Consequently, certain locally salient concerns or opportunities may be underrepresented. Future work would benefit from integrating participatory or interview-based approaches to broaden the evidence base and improve the relevance of ecosystem service and disservice assessments for decision-making. Such extensions were beyond the scope of the present study due to time and resource constraints.

## Conclusions

Megaherbivore rewilding may represent a relevant restoration strategy for traditional rural wood-pastures if stakeholders recognize the role of herbivory disturbance in sustaining the ecological dynamics, cultural heritage, and semi-open habitat mosaics that have historically characterized these systems (Garrido *et al*., 2021; Trouwborst, 2025). Our results suggest that its potential is strongest where reintroductions would expand pivotal ecological effects, such as restoring grazing-dominated disturbance regimes, contributing to fire-risk mitigation, and supporting nature-based rural economies. However, these benefits are not automatic. Rewilding may also generate socioecological disservices, including excessive herbivory pressure on conservation-priority habitats, crop and forestry damage, disease transmission, road collisions, management costs, and conflicts among stakeholders. Notably, effective management likely require population-density regulation, which could itself become socially contentious or be opposed by some conservation actors, while perceived rewilding risks may increase public fear of wildlife, particularly in relation to potential disease transmission (Thulin *et al*., 2015). These issues should therefore be explicitly addressed during the preparatory phase of any rewilding intervention (Carver *et al*., 2021). Future research should move beyond broad assessments of rewilding potential and develop spatially explicit, density-dependent, and context-specific scenarios. Such assessments will be essential to identify where megaherbivore rewilding can enhance traditional wood-pasture systems conservation without transferring disproportionate costs to rural communities.

